# Less is more: Coarse-grained integrative modeling of large biomolecular assemblies with HADDOCK

**DOI:** 10.1101/715268

**Authors:** Jorge Roel-Touris, Charleen G. Don, Rodrigo V. Honorato, João P.G.L.M Rodrigues, Alexandre M.J.J. Bonvin

**Affiliations:** Bijvoet Center for Biomolecular Research, Faculty of Science – Chemistry, Utrecht University. The Netherlands; Department of Pharmaceutical Sciences, University of Basel. Switzerland; Department of Structural Biology, Stanford University School of Medicine, CA. USA

**Keywords:** HADDOCK, protein-protein docking, MARTINI, coarse-graining, integrative modelling.

## Abstract

Predicting the 3D structure of protein interactions remains a challenge in the field of computational structural biology. This is in part due to difficulties in sampling the complex energy landscape of multiple interacting flexible polypeptide chains. Coarse-graining approaches, which reduce the number of degrees of freedom of the system, help address this limitation by smoothing the energy landscape, allowing an easier identification of the global energy minimum. They also accelerate the calculations, allowing to model larger assemblies. Here, we present the implementation of the MARTINI coarse-grained force field for proteins into HADDOCK, our integrative modelling platform. Docking and refinement are performed at the coarse-grained level and the resulting models are then converted back to atomistic resolution through a distance restraints-guided morphing procedure. Our protocol, tested on the largest complexes of the protein docking benchmark 5, shows an overall ~7-fold speed increase compared to standard all-atom calculations, while maintaining a similar accuracy and yielding substantially more near-native solutions. To showcase the potential of our method, we performed simultaneous 7 body docking to model the 1:6 KaiC-KaiB complex, integrating mutagenesis and hydrogen/deuterium exchange data from mass spectrometry with symmetry restraints, and validated the resulting models against a recently published cryo-EM structure.

## INTRODUCTION

Proteins are the workhorses of the cellular machinery. In order to function, they bind to one another, as well as to other biomolecules, to form large molecular assemblies. These interactions play a key role in all essential molecular processes within a cell. Most of these assemblies may exist as transient associations, which, together with other experimental factors, makes the characterization of their three dimensional (3D) structure a challenge^1^ for experimental methods such as nuclear magnetic resonance (NMR) spectroscopy or X-ray crystallography^2,3^. Despite recent advances in cryo-electron microscopy (cryo-EM), it is unlikely that the substantial gap between the number of estimated protein-protein interactions and those deposited in the Protein Data Bank^4^ can be overcome based solely on experimental methods^5^.

Computational docking has come of age as a complement to experimental methods in order to generate 3D models of protein assemblies. In particular, data- or information-driven docking and other integrative approaches are particularly appealing^1,6–8^. While docking performs sufficiently well for small- and medium-sized proteins, applications to large biological systems, either containing large individual molecules or a large number of interactors, are limited by the significant computational cost of thoroughly sampling complex conformational landscapes. Coarse-grained (CG) models mitigate this limitation by grouping atoms into larger “pseudo-atoms” or beads^9–11^, thus reducing the number of particles to consider in the computations. These models were used in the very first energy minimization of a protein in 1969^12^ and again in the first docking simulation^13^.

Since then, several CG models have been developed and applied to study different aspects of protein structural biology^14^. For protein docking in particular, of the CG models developed over the years, three standouts for their performance and/or success in community assessment experiments: Those implemented in ATTRACT, CABS-dock, and RosettaDock. The ATTRACT model^15,16^, developed by Zacharias and coworkers for flexible protein docking, represents the protein backbone by two pseudo-atoms and the side chains by an additional particle (or two in the case of larger amino acids). Non-bonded interactions are described by 8-6 LJ potentials and a Coulomb type term^17^, with parameters systematically optimized on both existing structures of protein-protein complexes as well as on docked models. As such, this limits the transferability of ATTRACT to other systems, such as protein-nucleic acid complexes or membrane proteins. Another model, CABS (Cα-Cβ-Side group protein model) was originally developed for structure prediction of globular proteins^18^ and later applied to protein-peptide docking^19^ (CABS-dock). As in ATTRACT, protein residues are represented by a maximum of four particles: Cα, Cβ, side chain and an extra particle representing a virtual Cα-Cα bond. Knowledge-based statistical potentials are used to describe particle interactions. The performance of CABS-dock was benchmarked on a set of protein-peptide complexes^20^, with peptides of 5-15 residues in length yielding accurate predictions. Although there are no technical limitations to the application of CABS-dock to larger protein-protein systems, except the increase in computational time, this application has not been reported in the literature to date and its performance remains thus uncertain. Moreover, given the specificity of its parameters to proteins, much like ATTRACT, the transferability of CABS to other molecular systems might be limited.

Finally, RosettaDock implements a two-step protocol with a coarse-grained global search followed by an all-atom refinement^21^. In the coarse-grained step, the interacting proteins are represented by their backbone atoms and a single pseudo-atom for the side chain. The resulting models are ranked using a combination of residue pairwise interaction terms, a contact-based term, and a term that penalizes overlapping residues. The all-atom refinement step uses the full Rosetta scoring function. As such, in the case of large assemblies, RosettaDock benefits from a smoother energy landscape during the conformational sampling but the second all-atom refinement stage is computationally expensive.

On the other hand, some CG models were developed to be easily transferrable. MARTINI, a CG model for biomolecules, was originally applied to study lipid bilayer assembly^22^ and later extended to proteins^23^, carbohydrates^24^ and nucleic acids^25,26^. This model maps, generally, four heavy atoms onto one coarse-grained bead. Its corresponding force field parameters have been calibrated to reproduce thermodynamic measurements. Systems are represented by 4 different basic particle types - polar (P), nonpolar (N), a-polar (C) and charged (Q) – that are further divided based on their hydrogen-bonding properties and their degree of polarity, giving a total of 18 unique “building blocks”. In addition to the 4 standard types of beads, the 2.2p version of MARTINI includes off-center charges for polar and charged amino acids. These extra “fake beads” improve the description of interactions between charged residues (ARG, LYS, ASP, GLU), and provide directionality/orientation in the case of polar residues, mimicking to some extent hydrogen bonds (e.g. an ASN side-chain bead has two “fake beads” associated carrying a small positive and negative charge, respectively). In addition, the MARTINI model is able to represent several types of molecules and allows for a straightforward conversion to atomistic resolution, making it ideal to use in HADDOCK for integrative modeling applications.

Here, we describe the implementation of the MARTINI CG force field for proteins^27^ in our information-driven docking software HADDOCK^28^. We evaluated the performance of the coarse-grained HADDOCK protocol using the largest complexes from the protein docking benchmark 5^29^, comparing it to the standard all-atom protocol. The performance increase from using a smaller set of particles to describe the molecular system allows for a substantial decrease in computational time, enabling the modeling of larger systems. As a demonstration, we modelled the heptameric KaiC-KaiB 1:6 assembly, which is part of the endogenous biological clock in cyanobacteria^30,31^, by performing a simultaneous 7 body docking, guided by mass-spectrometry (MS) and mutagenesis data in combination with symmetry restraints.

## METHODS

### Implementation of MARTINI in HADDOCK

The integration of the MARTINI CG force field for proteins into HADDOCK focused on three key aspects: (1) Converting the topology description and parametrization for each amino acid in a format suited for HADDOCK and its computational engine CNS (Crystallography and NMR System^33,34^), (2) adapting the atomic solvation parameters^35^ used to calculate the desolvation energy in HADDOCK to the CG particles and, (3) developing a protocol to convert the coarse-grained system back to atomistic resolution after the semi-flexible refinement stage of HADDOCK, making use of distance restraints derived from the MARTINI atoms-to-bead mapping.

As in standard MARTINI, four types of interaction sites are considered: Polar (P), non-polar (N), a-polar (C) and charged (Q). The conversion of the backbone to the CG beads follows a four-to-one (4:1) mapping rule, where all four heavy atoms (N, C_α_, C, O) are represented by a single bead placed at their geometric center. The conversion of side-chains varies, ranging from the same 4:1 mapping to 2:1 mapping and “small” beads in rings (HIS, PHE, TYR, TRP). We converted the topology and corresponding parameters of MARTINI 2.2p to a format compatible with CNS (see *Tables SI-1-4* in *Supplementary Information*). The force field, however, does not account for either the various possible histidine charge states (i.e. neutral with the proton on either the δ or ε nitrogen atom, or doubly-protonated and positively charged) nor for nonstandard residues (e.g. amino acids with post-translational modifications) or cofactors.

Since the amino acid backbone parameters in MARTINI are secondary structure-dependent, we use DSSP^36,37^ to analyze the initial structures and encode the secondary structure in the B-factor field. Using the later information HADDOCK automatically selects the proper parameters for each backbone bead in the coarse-grained structures when building the topology of the system. This effectively restrains the existing secondary structures, which might be a limitation for docking cases with large conformational changes between the unbound and bound states. However, if no secondary structure information is encoded in the B-factor field, random coil parameters allowing for possible conformational changes will apply. Note that in contrast to standard molecular dynamics simulations of proteins using the MARTINI force field, no Go terms are used in HADDOCK since only the interface is refined and therefore the majority of the structure is kept rigid by default.

Non-bonded CG interactions are calculated using a 14Å cutoff, as recommended, while interactions between atoms in the final stage are calculated using the OPLS force field^38^ parameters with the default 8.5Å cutoff used in HADDOCK.

### Solvation parameters for the coarse-grained particles

The HADDOCK score, used to rank the predicted models, is a linear combination of energetical and empirical terms (see *Scoring* below), including a solvent-accessible surface-based desolvation energy term^35^ (E_desolv_). In order to score CG models using this desolvation energy, we mapped the atomistic solvation parameters onto the CG beads. For this, the solvation energy of each group of atoms belonging to a specific bead was calculated for all 20 amino acids *X* in a *GGXGG* peptide. Since the solvation energy depends on the solvent accessible surface area of an atom/bead, the total atomistic energy was divided by the solvent accessible surface area of the corresponding CG bead in a similar peptide in order to obtain the CG solvation parameters *SP^i^*_*cg*_ for a specific CG particle *i* (Eq. 1):

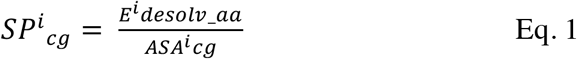

where *E*^*i*^_*desolv_aa*_ is the atomistic solvation energy for the group of atoms belonging to a given bead *i* and ASA^i^_cg_ is the accessible surface area of that bead in the *GGXGG* peptide.

The all-atom and CG solvent accessible areas were calculated using CNS with an accuracy of 0.0025 using a water radius of 1.4Å excluding all hydrogen atoms. The so-called *“fake beads”* are not included in the desolvation energy calculation. The resulting solvation parameters values for the MARTINI CG beads are listed in *Table 1*.

**Table 1.** Coarse-grained solvation parameters for each amino acid, mapped from the all-atom empirical solvation parameters onto MARTINI beads. BB: backbone beads. SC*: any side-chain bead. Note that “*fake beads*” (SCD) are not considered.

| Amino Acid | Solvation Parameter<br>$\frac{Kcal}{mol * \text{\AA}^2}$ | |
| --- | --- | --- |
|  | BB | SC* |
| ALA | -0.0107 | - |
| GLY | -0.0089 | - |
| ILE | -0.0153 | 0.0255 |
| LEU | -0.0153 | 0.0243 |
| VAL | -0.0158 | 0.0222 |
| PRO | -0.0046 | 0.0230 |
| ASN | -0.0137 | -0.0192 |
| GLN | -0.0147 | -0.0135 |
| THR | -0.0165 | -0.0009 |
| SER | -0.0154 | -0.0056 |
| MET | -0.0130 | 0.0202 |
| CYS | -0.0167 | 0.0201 |
| PHE | -0.0126 | 0.1005 |
| TYR | -0.0134 | 0.0669 |
| TRP | -0.0134 | 0.0872 |
| ASP | -0.0169 | -0.0360 |
| GLU | -0.0150 | -0.0301 |
| HIS | -0.0155 | 0.0501 |
| LYS | -0.0163 | -0.0210 |
| ARG | -0.0162 | -0.0229 |

### Preprocessing of input structures for coarse-grained docking

Setting up a CG docking run requires first converting the coordinate files, which contain information on individual atoms, into a CG representation. To this end, we adapted the “*martinize1.1py*” (https://github.com/Tsjerk) to account for the name type extensions (i.e. *“fake beads”* present in the 2.2p version of MARTINI) and to additionally generate distance restraints, in CNS format, between the original atoms and the corresponding CG beads, which are used in the final back-mapping stage of the protocol (see below). Since the MARTINI backbone parametrization depends on the local secondary structure, we numerically store the secondary structure assignments computed by DSSP^36,37^ into the B-factor column of the resulting CG PDB files. As in the standard protocol, HADDOCK automatically builds any missing atom when creating both the topology and coordinate files from the user-provided PDB files. This procedure is done both for the starting CG and all-atom structures. The latter are used in the final back-mapping stage from CG to all-atom.

### Back-mapping coarse-grained models to atomic resolution by distance restraints

In order to convert the final coarse-grained models back to an all-atom representation, we make use of the ability of HADDOCK to use distance restraints to guide the modelling, using the atom-to-bead distance restraints derived during the initial setup stage. For a group of atoms belonging to a particular CG bead, we create one distance restraint with 0 length between the geometric center of the atoms and the bead to which they belong. The conversion protocol consists of the following steps:

#### (1) Initial fitting onto the CG model

The all-atom structure of each molecule of the complex is fitted onto its respective CG representation in the docked CG model by rigid body energy minimization (EM) guided by the CG-to-AA distance restraints. During this step the CG model is kept fixed and the intermolecular interactions are scaled by a factor 0.001 to account for possible clashes between the AA molecules. No energy terms are included for the CG model, except the distance restraining potential.

#### (2) Inducing conformational changes

In order to morph the all-atom structure onto the CG model, which might have undergone conformational changes during the flexible stage of the docking protocol, we first perform two short rounds of energy minimization (500 steps), increasing the scaling factor for intermolecular interactions to 0.01 after the first minimization. Then, we perform 500 steps of Cartesian molecular dynamics (MD) at 300K with an integration time step of 0.0005 ps and another round of EM.

#### (3) Clearing clashes and optimizing all-atom interactions

We perform two rounds of energy minimization, increasing the scaling factor of the intermolecular interactions to 0.1 and 1.0, respectively, followed by another short MD (500 integration steps) and two extra minimization rounds.

In all three steps, all covalent and non-covalent energy terms are included for the AA models together with the restraint energy term for the atom-to-bead distance restraints. Once the all-atom models have been generated, the CG models are discarded, the morphing distance restraints are removed and all other restraining energy terms representing the various data given to HADDOCK to drive the docking are reintroduced. These are used in a final round of energy minimization. Although computational expensive for large systems, the user can then choose to follow-up with the full water refinement stage of the standard HADDOCK protocol (turned off by default).

### Docking procedure

All docking calculations were performed using a local installation of the new HADDOCK version 2.4 supporting CG docking. This protocol is also supported by the new version of our webserver^39^ soon to be released. For comparison purposes, the docking was performed both with all-atom and coarse-grained representations, using the united-atom OPLS force-field^40^ and MARTINI 2.2p, respectively. The docking was guided by ambiguous interaction restraints (AIRs) derived from the bound complexes (true interface) by selecting all solvent accessible residues with at least one heavy atom within 3.9Å from any heavy atom of the partner molecule. These restraints represent an ideal scenario where accurate information is available about the residues in the interface but not about their specific pairwise contacts (information that can be obtained e.g. from NMR chemical shift perturbations, mass-spectrometry hydrogen/deuterium exchange, …)^7,8^. The sampling parameters were kept as default in HADDOCK: 1000/200/200 models were generated for the rigid body (it0), semi-flexible (it1), and water refinement (itw) stages, respectively. In the CG runs, the final water refinement stage was replaced by the back-mapping from CG to all-atom as shown in *Fig.1*. The final models were clustered based on the fraction of common contacts (FCC)^41^ using a 0.6 cutoff and a minimum number of 4 models per cluster.

**Fig 1.**
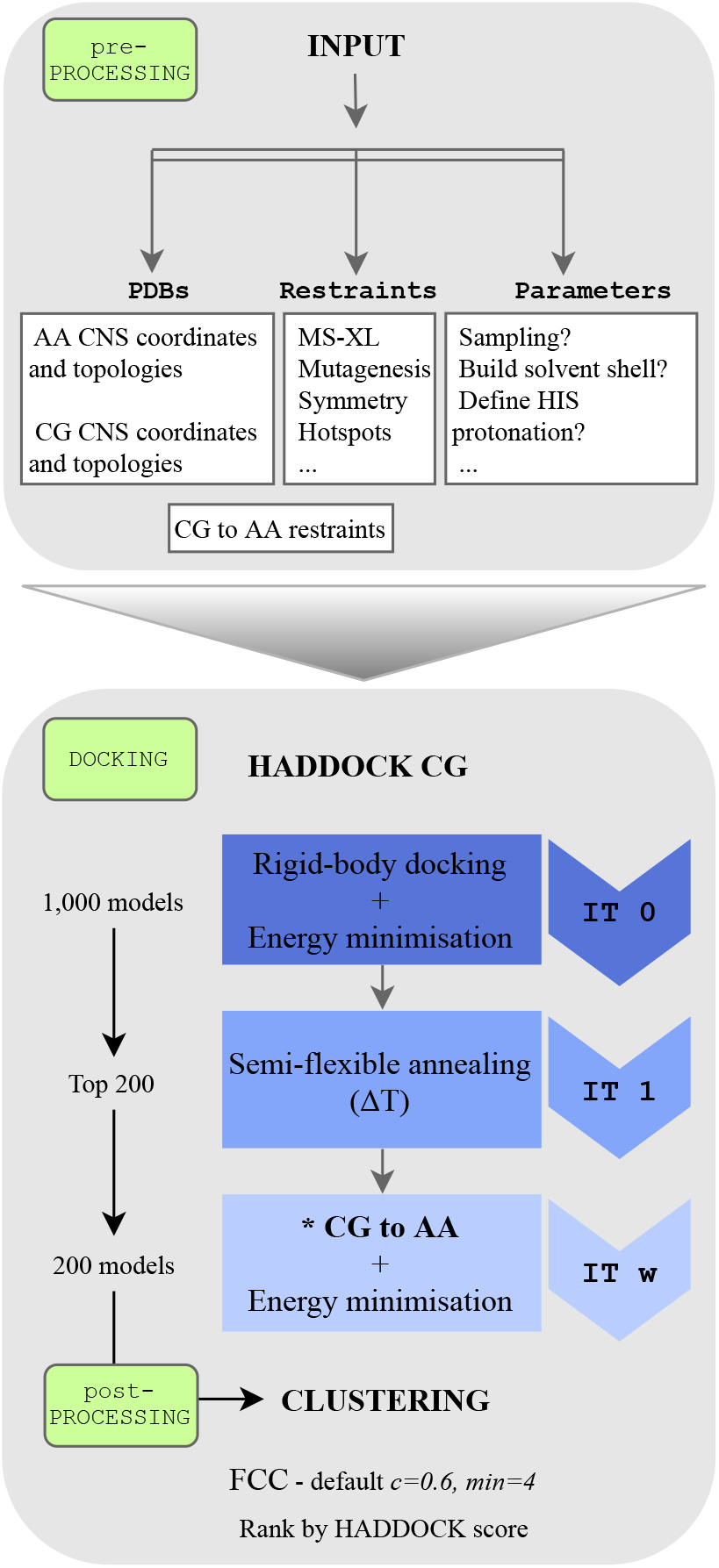
HADDOCK coarse-grained flowchart. Default protein-protein coarse-grained protocol in HADDOCK. AA = all-atom, CG = coarse-grained, FCC = fraction of common contacts. * Back-mapping coarse-grained models to atomic resolution by distance restraints.

### Scoring

We investigated whether re-parameterizing the HADDOCK-CG score led to a better scoring performance by systematically varying the weights of the scoring function. Since we did not observe significant improvements (data not shown), we kept the original HADDOCK scoring functions (HS) for the three stages of the docking protocol (rigid-body EM (it0); semi-flexible refinement (it1); explicit solvent refinement (itw)):

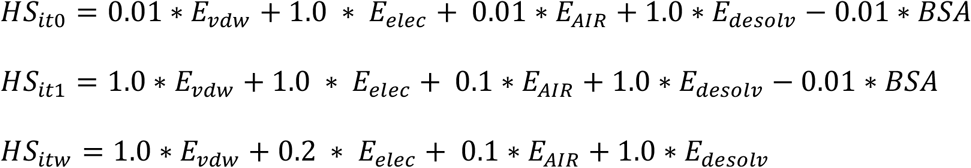

where E_vdw_ and E_elec_ are the van der Waals and electrostatic energies terms calculated using a 12-6 Lennard-Jones and Coulomb potential, respectively, with MARTINI (it0, it1) or OPLS (itw) non-bonded parameters, E_AIR_ is the ambiguous interaction restraints energy, E_desolv_ is the empirical desolvation score and BSA is the buried surface area in Å^2^.

### Protein docking benchmark

To test the performance of our HADDOCK-CG protocol, we selected a subset of complexes from the Protein-Protein Docking Benchmark version 5.0^29^, consisting of all complexes with more than 5,000 heavy atoms, excluding all antibody-antigen cases. This selection yielded a benchmark set of 27 cases (see *Table SI-5* in *Supplementary Information*).

### Metrics for evaluation of model quality

The quality of the generated models was evaluated using standard CAPRI^42^ criteria, including the fraction of native contacts (FNAT) and the positional interface (i-RMSD) and ligand (l-RMSD) root mean square deviations from the reference crystal structure. FNAT is calculated using all heavy atom – heavy atom intermolecular contacts using a 5Å distance cutoff (CAPRI definition)^42^. The i-RMSD is calculated on the interface after superimposition on the interface residues, defined as those with any heavy atom within a 10Å distance of the partner protein. The l-RMSD is calculated on the ligand (usually the smallest molecule) after superimposition on the backbone atoms of the receptor (largest molecule). For both, i-RMSD and l-RMSD, only backbone heavy atoms are considered (C_α_, C, N, O). Based on these three metrics, the quality of the docking poses is classified as:

- High: FNAT ≥ 0.5 and i-RMSD ≤ 1Å or l-RMSD ≤ 1Å,
- Medium: FNAT ≥ 0.3 and 1Å < i-RMSD ≤ 2Å or 1Å < l-RMSD ≤ 5Å,
- Acceptable: FNAT ≥ 0.1 and 2Å < i-RMSD ≤ 4Å or 5Å < l-RMSD ≤ 10Å,
- Near-Acceptable: FNAT ≥ 0.1 and 4Å < i-RMSD ≤ 6Å and
- Low quality: FNAT < 0.1 or i-RMSD > 6Å or l-RMSD > 10Å.

### Metrics for the evaluation of the docking success rate

The performance of the docking calculations was analyzed as follows: (1) The percentage of cases in which at least one model of a given accuracy is found within the top *N* solutions ranked by HADDOCK (*N* = 1, 5, 10, 20, 25, 50, 100, 200), and (2) the percentage of cases in which at least one acceptable or higher quality model was found in the top *T* clusters (*T* = 1, 2, 3, 4, 5).

### KaiC-KaiB coarse-grained integrative modelling with HADDOCK

In order to model the KaiC:KaiB 1:6 complex we performed two different docking runs, targeting either the CI or CII domains on KaiC since the H/D exchange data from MS point to two possible interfaces (for details refer to Snijder et al^43^). We used the crystal structure of KaiC (PDB ID: *3DVL*) consisting of 12 domains (two 6-membered rings) as starting point for the docking. For KaiB, we used six copies of the recent NMR structure (PDB ID: *5JYT*)^44^, which shows a fold-switch at the interacting region compared to the previously-determined crystal structure^45^.

The regions experimentally identified by HDX-MS as protected from solvent in either the CI or CII domains of KaiC and in KaiB were specified as active residues in HADDOCK, after filtering them for solvent accessibility (relative residue solvent accessibility larger than 50% as calculated with NACCESS^46^) (see *Table SI-6* in *Supplementary Information*, for a detailed list of residues). For KaiB, we included three additional residues identified by mutagenesis experiments. A structural similarity analysis of KaiC revealed an asymmetrical structure with RMSD values for the interface regions between subunits in the hexamer ranging from 0.9Å to 1.9Å (see *Table SI-7* in *Supplementary Information* for more details). As a result, we restrained the KaiB monomers to an approximate C6 symmetry by defining three C2 symmetry pairs (B-E/C-F/D-G) and two C3 symmetry triplets (B-D-F/C-E-G), but we did not use non-crystallographic symmetry restraints (NCS) since the interfaces are asymmetrical.

Because of the symmetry restraints, sampling of 180º rotations during the rigid-body stage was disabled. Furthermore, given the large size of the complex and the number of subunits to dock (7), the sampling was increased to 10000/400/400 models for it0/it1/itw, respectively. Finally, we disabled the final refinement in explicit water, only performing the back-mapping from CG to all-atom (as part of the default HADDOCK-CG pipeline). We only used the top 200 models according to the HADDOCK score for analysis and validation purposes.

## RESULTS AND DISCUSSION

We have integrated the MARTINI 2.2p force field for proteins into HADDOCK (see Methods), adapted the desolvation energy terms to the coarse-grained beads, and developed a distance restraints-based back-mapping method to restore the atomic resolution of the final models while accounting for possible conformational changes that took place during the CG semi-flexible refinement step. In the following sections, we discuss the performance of our protocol in terms of success rate, sampling and computational efficiency using the 27 largest complexes from the docking benchmark 5. We then showcase its potential by modelling a large heptameric complex using mass-spectrometry and mutagenesis data.

### Overall performance of coarse-grained HADDOCK

We compared the unbound docking performance of HADDOCK-CG with the default all-atom protocol for 27 binary complexes from the docking benchmark 5 (see Methods; *Protein docking benchmark*). Fourteen of those complexes were classified as easy according to the structural differences between the bound and unbound structures of the monomers, 8 as medium and 5 as hard. The docking was performed starting from the unbound structures of each protein and driven by information from the real interface (see Methods; *Docking* procedure), mimicking an ideal scenario for HADDOCK users. The success rate was defined as the percentage of cases for which an acceptable or better model was obtained in the top *N* ranked models (for details see Methods; *Metrics for evaluation of success in docking*).

Coarse-grained docking shows a slightly better overall performance (*Fig. 2*) in the top 1 for single structure ranking (best ranked structure) than the standard all-atom protocol, with success rates for acceptable or higher quality models of 51.8% and 48.1% respectively. However, this trend reverses for the performance in the top 5, with 66.6% and 77.7% success rates for coarse-grained and atomistic models, respectively. For the remaining top *N*, (*N* = 10, 20, 50, 100, 200), the performance of HADDOCK-CG is comparable with that of all-atom calculations, reaching a maximum of 92.5% at N = 200. For the two cases with the largest conformational change (i-RMSD values of 4.69Å / 5.79Å between unbound and bound structures for *1Y64* / *4GAM, respectively*), neither coarse-grained nor all-atom calculations generated near-acceptable solutions.

**Fig 2.**
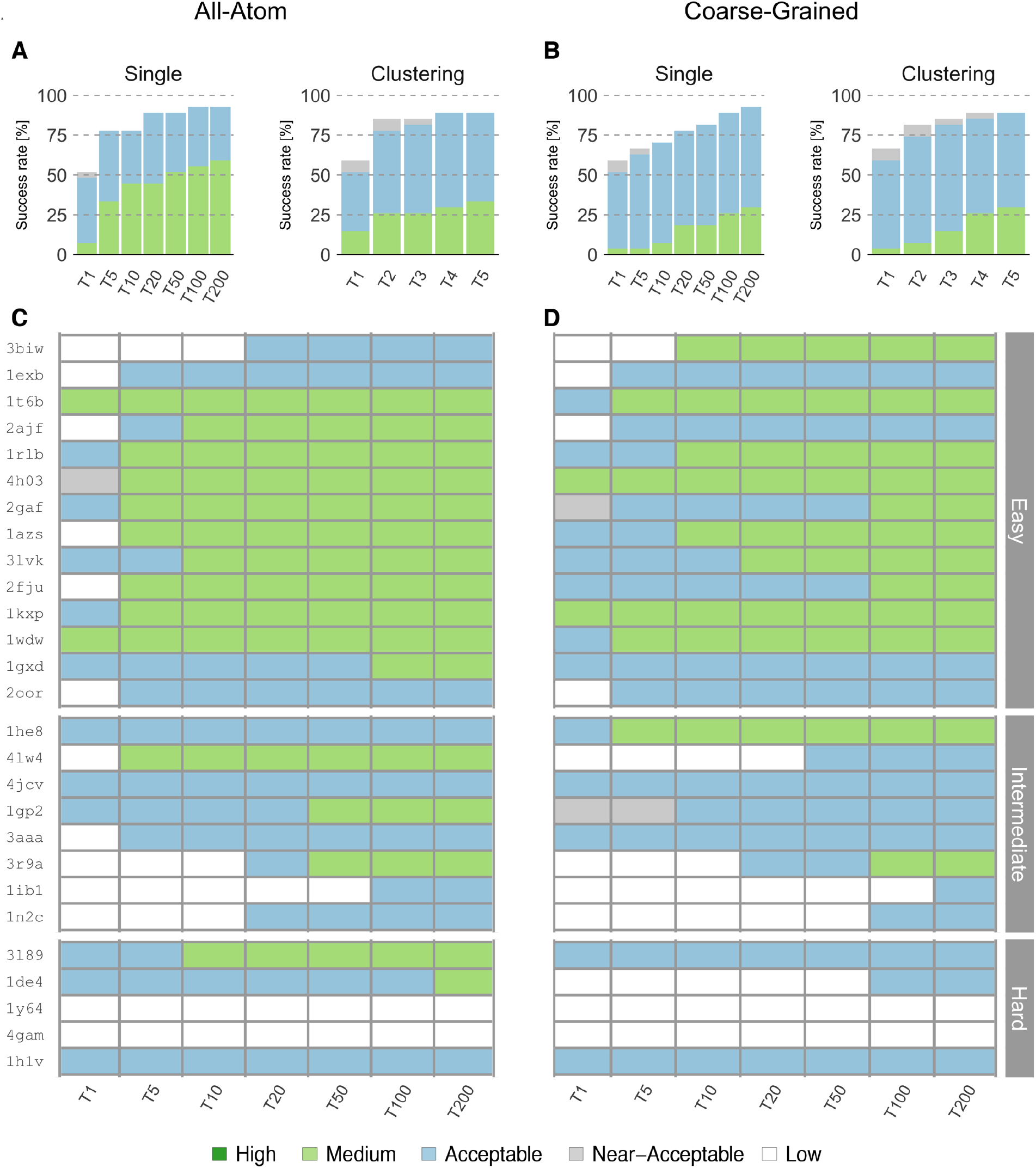
Performance of the all-atom and coarse-grained protocols in HADDOCK on the 27 largest complexes of the Docking Benchmark 5. (**A**) Overall success rates (%) of the all-atom protocol on ranking single models (Single) or clusters (Clustering) as a function of the number of models/clusters considered. (**B**) Same as (A) but for the coarse-grained protocol. (**C**) and (**D**), Quality of the docking models for all 27 cases as a function of the number of models considered. The complexes are ordered by increasing degree of difficulty (from top to bottom) for both all-atom and CG docking runs. The color coding indicates the quality of the docked models.

We also analyzed the success rate on a per-cluster basis, which is the standard scoring scheme of HADDOCK. Clustering models improves the success rate for both coarse-grained and all-atom simulations to 59.2% and 51.8%, respectively, for the top 1 cluster. The success rate is maximal for the top 5 clusters reaching 88.8% for acceptable or higher quality models (*Fig. 2B*). The all-atom protocol reached the maximum success rate (88.8%) at the top 4 clusters. Compared to single structure scoring, no near-native cluster was obtained for *1IB1* due to the fact that only 3 models passed the quality thresholds and our clustering strategy requires a minimum of 4 models per cluster.

Concerning the quality of the models (see Methods; *Metrics for evaluation of model quality*), the all-atom runs generated higher quality solutions than CG runs (*Fig. 2C and 2D*). For the easy cases, all-atom runs rank medium quality models in the top 10 solutions for 10 out of 14 cases and acceptable quality models for 13 out of 14 cases. For the CG runs, medium quality models are obtained in the top 10 solutions for 7 out of 14 easy cases, and acceptable quality models for all 14 cases. As for the intermediate and hard cases, the all-atom runs generate medium quality models for only 5 out of 13 cases, while CG runs generate them in 2 cases. Overall, coarse-grained HADDOCK generated medium quality solutions for 12 out of all 27 complexes including intermediate cases, slightly worse than the 16 cases for the all-atom runs.

Interestingly there are 2 cases where CG docking generates better quality models than all-atom runs. For *3BIW*, an easy case, coarse-grained docking generated medium quality models ranked in the top 10. The best of these models has an FNAT of 0.61 and i-RMSD of 1.9Å, compared to FNAT of 0.52 and i-RMSD of 3.5Å for the all-atom run. For *1HE8*, a medium difficulty case, we found a medium quality model in the top 5 with an FNAT of 0.55 and l-RMSD of 4.9Å, while the best all-atom model has an FNAT of 0.44 and l-RMSD of 6.1Å.

Given the back-mapping to all-atom resolution at the end of the coarse-grained protocol, we also evaluated the quality of the final models in terms of the number of atomic clashes at the interface. A clash was defined as any pair of heavy atoms belonging to different molecules within 3Å distance, in accordance with the CAPRI assessment procedure^47^. The number of clashes was then divided by the buried surface area of the complex and models with more than 0.1 clashes/Å^2^ were considered of poor quality. We found no model, in both CG and all-atom runs, that scored under this clash threshold. However, and interestingly, docked structures generated via coarse-graining presented, on average, half the clashes of the models from the all-atom runs, which might be explained by the multiple energy minimization rounds performed during the back-mapping protocol, compared to the default water refinement protocol.

### Reduction of the energy landscape complexity

A product of coarse-graining is a smoothening of the energy landscape, which should allow for an easier sampling compared to all-atom calculations. The coarse-grained landscape might help find energy minima, especially in cases where only few or no data are available to drive the modelling and should, therefore, contribute to a better performance of coarse-grained docking runs (i.e. an increase in the number of near-acceptable models). To test this hypothesis, we performed docking without any experimental information, using the ab initio mode of HADDOCK in which, for each docked model, pairs of residues on the interacting molecules are randomly selected and ambiguous interaction restraints are defined between surface patches within 7.5Å of those residues. In order to test whether coarse-graining improves sampling, we ran our benchmark with this type of random restraints for both all-atom and coarse-grained protocols, increasing in both case the sampling to 10000/400/400 models for it0/it1/itw. We indeed observe (*Table 2*) a substantial increase (28.4%) in the number of models of acceptable or better quality during the rigid body stage of coarse-grained docking, compared to all-atom simulations. However, when using interface data to drive the calculations, this difference decreases to 8% more acceptable or higher quality models for the coarse-grained protocol, which is still a substantial improvement.

**Table 2.** Comparison of the total number of acceptable or higher quality models, generated over all 27 complexes at the rigid-body stage (it0), between coarse-grained and standard all-atom HADDOCK protocols in the absence of information to drive the docking (*ab-initio* mode) and using true interface information. 10000 models were generated in the case of abinitio docking. For details, see *Tables SI-10-11* in *Supplementary Information*.

| *Ab-initio docking (random patches)* |  |  |  |  |
| --- | --- | --- | --- | --- |
| Coarse-grained | 15 | 16 | 74 | **1.39** |
| All-atom | 11 | 13 | 53 |  |
| *True interface docking* |  |  |  |  |
| Coarse-grained | 2666 | 5066 | 9689 | **1.08** |
| All-atom | 2702 | 4940 | 8896 |  |

### Computational performance

The main motivation to implement a coarse-grained forcefield in HADDOCK was to accelerate and enable the modelling of large biomolecular assemblies by reducing the number of particles considered during the computations. The atom-to-bead mapping of the MARTINI model leads to a significant reduction in the number of particles, making the computations substantially more efficient. It was previously shown that MARTINI allows for an increase in computational efficiency by a factor 2 to 4 compared to common all-atom models^23^. In our case, integrating MARTINI into HADDOCK led to an average ~7-fold speed-up in total computation time (*Table 3*).

**Table 3.**
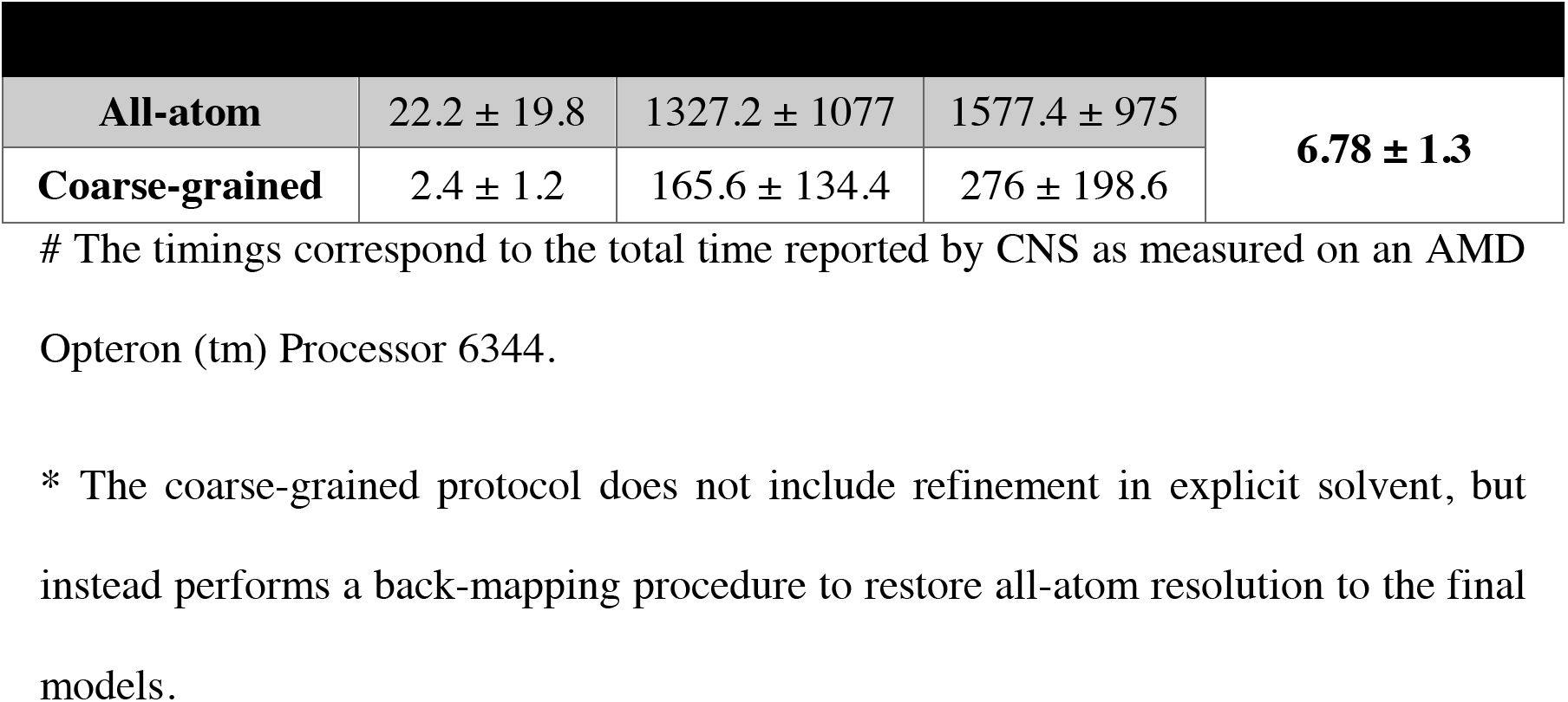
Comparison of average CPU times (seconds/model)^#^ for the test benchmark (N = 27) between the all-atom and coarse-grained HADDOCK protocols.

### Coarse-grained integrative modelling of KaiC-KaiB

To demonstrate our coarse-grained HADDOCK protocol, we modelled the heptameric KaiC-KaiB (stoichiometry1:6) complex by simultaneous 7 body docking using data from mutagenesis experiments and hydrogen-deuterium exchange MS^43^. The structures of KaiC and KaiB have been both characterized individually at atomic level. KaiC forms hexamers and consists of two domains, CI and CII^48,49^. It has been shown that six KaiB monomers bind to one KaiC hexamer^30^. The first published model of this complex^43^ wrongly pointed to CII as binding mode, based on better agreement with collision cross section data obtained by time of flight MS. Later on, the cryo-EM structure^50^ of KaiCBA revealed a CI binding mode and a different fold of KaiB corresponding to the solution NMR structure (PDB ID *5JYT*) that was solved after the initial model was published. This NMR structure, which is also the conformation found in the cryo-EM structure, shows a fold switch compared to the crystal structure (PDB ID *4KSO*) that was used in the initial modelling. The crystal structure was the only available one at the time of the first modelling. The first model was built by docking one KaiB onto two domains of KaiC (out of the 12 domains in full KaiC). We repeated here this modelling, using this time the full KaiC structure and six copies the binding competent KaiB conformation (the NMR structure). Two 7 body docking runs targeting the CI and CII binding interfaces were performed with HADDOCK-CG. Along with the experimental data, we imposed symmetry restraints (C3 + C2, as an approximation of C6) between the 6 KaiB components. The resulting models were scored and ranked according to the HADDOCK score (see Methods; *Scoring*), including an additional energy term for the symmetry restraints. The cryo-EM map (*EMDB-3603*) was used for independent validation of the models.

Using the new, binding-competent KaiB structure we clearly identify the CI binding mode as the right answer, with a significantly lower HADDOCK score than CII: −216.7 ± 13.2 a.u. versus +44.5 ± 19 a.u. for the best cluster of each run (see *Table SI-8* in *Supplementary Information*). This model obtained based on mutagenesis and mass-spectrometry data is consistent with the recent cryo-EM model of the KaiC:KaiB:KaiA complex in a fully assembled state^50^ with a l-RMSD of 3.6Å, calculated over all six interfaces, for the best model of the top scoring cluster (for more details, see *Table SI-9* in *Supplementary Information*). We further validated our model by quantifying its agreement with the published cryo-EM map of the complex (*EMDB-3603*) using Chimera^51^: The correlation score of our model is 0.82, compared to 0.84 for the original cryo-EM backbone model (PDB ID *5N8Y*) as shown in *Fig. 3*.

**Fig 3.**
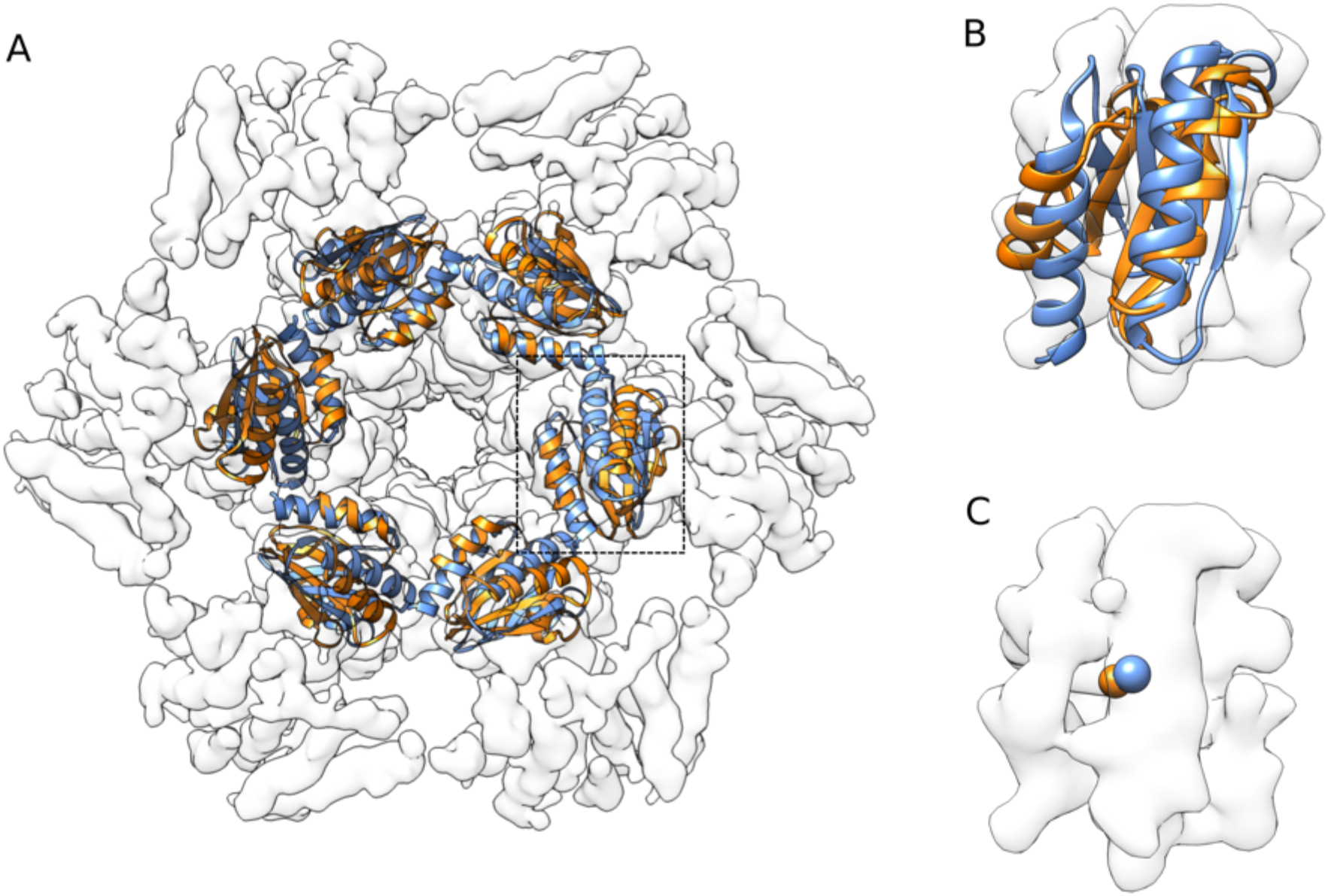
Comparison of the cryo-EM model (PDB code: 5N8Y, blue) and the best coarse-grained model obtained in this work (orange). The models were fitted into the map using Chimera^51^. The correlation coefficient for our docked model is 0.82 compared to 0.84 for the cryo-EM structure. **(A)** Top view of the KaiB hexamer bound to KaiC CI domain. **(B)** Detailed view of single KaiB. **(C)** Comparison of centers of mass of a single KaiB monomer. Note that KaiA present in the cryo-EM model is not shown here.

While the first all-atom model was obtained by docking a subset of the full complex, in this work we modelled here the full 1:6 KaiC-KaiB complex. By coarse-graining, we reduced the number of particles from 31726 in the original all-atom model to 9842 for the coarse-grained model, reducing the computational time by about a factor 6 times, from 4 hours to 48 minutes, on average, per model.

## CONCLUSIONS

In this work, we presented the integration of the MARTINI coarse-grained force field in our HADDOCK integrative modeling software. Our new docking protocol makes use of coarse-grained representations during the rigid body and semi-flexible refinement stages and restores the final docked models to atomistic resolution in a final back-mapping stage. By using distance restraints between beads and the atoms that belong to them, the back-mapping protocol is able to morph conformational changes that potentially took place during the coarse-grained flexible refinement. The performance of coarse-grained docking is similar to that of the standard all-atom protocol in terms of success rate and quality of the generated models. In addition, it generates more near-native models when limited or no data are available and comes with the benefit of a ~7-fold reduction in computing time. The power of our coarse-grained integrative modelling approach was demonstrated by modelling the structure of the heptameric KaiC:KaiB (1:6) complex, for which we obtained models in excellent agreement with the cryo-EM structure. In conclusion, the implementation of the MARTINI coarse-grained force field into HADDOCK extends its ability to model increasingly larger and more intricate biomolecular assemblies. In the future, we plan to make use of the MARTINI models for lipids and nucleic acids and extend our protocol to allow modelling of nucleic acid complexes, as well as membrane and membrane-associated complexes, for which we recently published a new docking benchmark^32^.

## ASSOCIATED CONTENT

### Supporting Information

The supporting information contains tables with all force field parameters converted to CNS format as well as an overview of the Docking Benchmark used. Furthermore, it includes a detailed list of residues used as restraints, paired i-RMSD values (KaiC starting structure), cluster-based statistics for the CI and CII docking runs and the structural similarity assessment of the top 4 models with respect to the cryo-EM data used for the integrative modelling of KaiCB. Finally, a per-complex based analysis of the number of acceptable or higher quality models generated at rigid-body (it0) stage of coarse-grained and standard all-atom protocols is reported.

## AUTHOR INFORMATION

### Author Contributions

Alexandre Bonvin and João Rodrigues designed the research with contributions of all authors. Charleen G. Don, João Rodrigues and Alexandre Bonvin performed the initial implementation into HADDOCK. Jorge Roel-Touris and Rodrigo V. Honorato implemented v2.2 of MARTINI and all the necessary machinery to convert AA to CG models.

Jorge Roel-Touris and Alexandre Bonvin performed the benchmarking and KaiC:KaiB modelling. All the authors contributed to the analysis of the data and the writing of the manuscript.

## ACKNOWLEDGMENT

The authors acknowledge all members from the Computational Structural Biology group at Utrecht University for fruitful discussions with special mention to Dr. Adrien Melquiond for his help on the integrative modelling of KaiCB. We thank the MARTINI group at Groningen University, the Netherlands, for their support in implementing MARTINI into HADDOCK. Finally, we acknowledge the use of software from the SBGRID consortium^52^ for various analysis tasks.

## ABBREVIATIONS

PPI: protein-protein interaction
NMR: nuclear magnetic resonance
AA: all-atom
CG: coarse-grain

## REFERENCES

(1) Ritchie, D. W. Recent Progress and Future Directions in Protein-Protein Docking. Curr. Protein Pept. Sci. 2008, 9 (1), 1–15.

(2) Aloy, P.; Pichaud, M.; Russell, R. B. Protein Complexes: Structure Prediction Challenges for the 21st Century. Current Opinion in Structural Biology. 2005, pp 15–22.

(3) Wass, M. N.; David, A.; Sternberg, M. J. E. Challenges for the Prediction of Macromolecular Interactions. Current Opinion in Structural Biology. 2011, pp 382–390.

(4) Berman, H. M.; Westbrook, J.; Feng, Z.; Gilliland, G.; Bhat, T. N.; Weissig, H.; Shindyalov, I. N.; Bourne, P. E. The Protein Data Bank. Nucleic Acids Res. 2000, 28 (1), 235–42.

(5) Mosca, R.; Céol, A.; Aloy, P. Interactome3D: Adding Structural Details to Protein Networks. Nat. Methods 2013, 10 (1), 47–53.

(6) Wiehe, K.; Peterson, M. W.; Pierce, B.; Mintseris, J.; Weng, Z. Protein-Protein Docking: Overview and Performance Analysis. Methods Mol. Biol. 2008, 413, 283–314.

(7) Karaca, E.; Bonvin, A. M. J. J. Advances in Integrative Modeling of Biomolecular Complexes. Methods. 2013, pp 372–381.

(8) Rodrigues, J. P. G. L. M.; Bonvin, A. M. J. J. Integrative Computational Modelling of Protein Interactions. FEBS J. 2014, 281 (8), 1988–2003.

(9) Rader, A. Coarse-Grained Models: Getting More with Less. Curr. Opin. Pharmacol. 2010, 10 (6), 753–759.

(10) Takada, S. Coarse-Grained Molecular Simulations of Large Biomolecules. Current Opinion in Structural Biology. 2012, pp 130–137.

(11) Saunders, M. G.; Voth, G. A. Coarse-Graining of Multiprotein Assemblies. Current Opinion in Structural Biology. 2012, pp 144–150.

(12) Levitt, M.; Lifson, S. Refinement of Protein Conformations Using a Macromolecular Energy Minimization Procedure. J. Mol. Biol. 1969, 46 (2), 269–279.

(13) Wodak, S. J.; Janin, J. Computer Analysis of Protein-Protein Interaction. J. Mol. Biol. 1978, 124 (2), 323–342.

(14) Kmiecik, S.; Gront, D.; Kolinski, M.; Wieteska, L.; Dawid, A. E.; Kolinski, A. Coarse-Grained Protein Models and Their Applications. Chemical Reviews. 2016, 116 (14), 7898–936.

(15) De Vries, S. J.; Schindler, C. E. M.; Chauvot De Beauchêne, I.; Zacharias, M. A Web Interface for Easy Flexible Protein-Protein Docking with ATTRACT. Biophys. J. 2015, 108 (3), 462–465.

(16) De Vries, S. J.; Rey, J.; Schindler, C. E. M.; Zacharias, M.; Tuffery, P. The PepATTRACT Web Server for Blind, Large-Scale Peptide-Protein Docking. Nucleic Acids Res. 2017, 45 (W1), W361–W364.

(17) Fiorucci, S.; Zacharias, M. Binding Site Prediction and Improved Scoring during Flexible Protein-Protein Docking with ATTRACT. Proteins Struct. Funct. Bioinforma. 2010, 78 (15), 3131–3139.

(18) Kolinski, A. Protein Modeling and Structure Prediction with a Reduced Representation. In Acta Biochimica Polonica; 2004, 51 (2), 349–71.

(19) Blaszczyk, M.; Kurcinski, M.; Kouza, M.; Wieteska, L.; Debinski, A.; Kolinski, A.; Kmiecik, S. Modeling of Protein-Peptide Interactions Using the CABS-Dock Web Server for Binding Site Search and Flexible Docking. Methods 2016, 93, 72–83.

(20) Kurcinski, M.; Jamroz, M.; Blaszczyk, M.; Kolinski, A.; Kmiecik, S. CABS-Dock Web Server for the Flexible Docking of Peptides to Proteins without Prior Knowledge of the Binding Site. Nucleic Acids Res. 2015, 43, W419–W424.

(21) Gray, J. J.; Moughon, S.; Wang, C.; Schueler-Furman, O.; Kuhlman, B.; Rohl, C. A.; Baker, D. Protein-Protein Docking with Simultaneous Optimization of Rigid-Body Displacement and Side-Chain Conformations. J. Mol. Biol. 2003, 331 (1), 281–99.

(22) Marrink, S. J.; Risselada, H. J.; Yefimov, S.; Tieleman, D. P.; De Vries, A. H. The MARTINI Force Field: Coarse Grained Model for Biomolecular Simulations. J. Phys. Chem. B 2007, 111 (27), 7812–7824.

(23) Monticelli, L.; Kandasamy, S. K.; Periole, X.; Larson, R. G.; Tieleman, D. P.; Marrink, S. J. The MARTINI Coarse-Grained Force Field: Extension to Proteins. J. Chem. Theory Comput. 2008, 4 (5), 819–834.

(24) López, C. A.; Rzepiela, A. J.; de Vries, A. H.; Dijkhuizen, L.; Hünenberger, P. H.; Marrink, S. J. Martini Coarse-Grained Force Field: Extension to Carbohydrates. J. Chem. Theory Comput. 2009, 5 (12), 3195–210.

(25) J. Uusitalo, J.; Ingólfsson, H. I.; Akhshi, P.; Tieleman, D. P.; Marrink, S. J. Martini Coarse-Grained Force Field: Extension to DNA. J. Chem. Theory Comput. 2015, 11, 3932–3945.

(26) Uusitalo, J. J.; Ingólfsson, H. I.; Marrink, S. J.; Faustino, I. Martini Coarse-Grained Force Field: Extension to RNA. Biophys. J. 2017, 113 (2), 246–256.

(27) De Jong, D. H.; Singh, G.; Bennett, W. F. D.; Arnarez, C.; Wassenaar, T. A.; Schäfer, L. V.; Periole, X.; Tieleman, D. P.; Marrink, S. J. Improved Parameters for the Martini Coarse-Grained Protein Force Field. J. Chem. Theory Comput. 2013, 9 (1), 687–697.

(28) Dominguez, C.; Boelens, R.; Bonvin, A. M. J. J. HADDOCK: A Protein-Protein Docking Approach Based on Biochemical or Biophysical Information. J. Am. Chem. Soc. 2003, 125 (7), 1731–1737.

(29) Vreven, T.; Moal, I. H.; Vangone, A.; Pierce, B. G.; Kastritis, P. L.; Torchala, M.; Chaleil, R.; Jiménez-García, B.; Bates, P. A.; Fernandez-Recio, J.; et al. Updates to the Integrated Protein-Protein Interaction Benchmarks: Docking Benchmark Version 5 and Affinity Benchmark Version 2. J. Mol. Biol. 2015, 427 (19), 3031–3041.

(30) Ishiura, M. Expression of a Gene Cluster *KaiABC* as a Circadian Feedback Process in Cyanobacteria. Science (80-.). 1998, 281 (5382), 1519–1523.

(31) O’Neill, J.; Reddy, A. Circadian Clocks in Human Red Blood Cells. Nature 2011, 469 (7331), 498–503.

(32) Koukos, P. I.; Faro, I.; van Noort, C. W.; Bonvin, A. M. J. J. A Membrane Protein Complex Benchmark. J. Mol. Biol. 2018, 430 (24), 5246–5256.

(33) Brünger, A. T.; Adams, P. D.; Clore, G. M.; Delano, W. L.; Gros, P.; Grossekunstleve, R. W.; Jiang, J. S.; Kuszewski, J.; Nilges, M.; Pannu, N. S.; et al. Crystallography & NMR System: A New Software Suite for Macromolecular Structure Determination. Acta Crystallogr. Sect. D Biol. Crystallogr. 1998, 54 (Pt5), 905–21.

(34) Brunger, A. T. Version 1.2 of the Crystallography and Nmr System. Nat. Protoc. 2007, 2 (11), 2728–33.

(35) Fernández-Recio, J.; Totrov, M.; Abagyan, R. Identification of Protein-Protein Interaction Sites from Docking Energy Landscapes. J. Mol. Biol. 2004, 335 (3), 843–865.

(36) Kabsch, W.; Sander, C. Dictionary of Protein Secondary Structure: Pattern Recognition of Hydrogen-bonded and Geometrical Features. Biopolymers 1983, 22 (12), 2577–2637.

(37) Touw, W. G.; Baakman, C.; Black, J.; Te Beek, T. A. H.; Krieger, E.; Joosten, R. P.; Vriend, G. A Series of PDB-Related Databanks for Everyday Needs. Nucleic Acids Res. 2015, 43, D364–D368.

(38) Jorgensen, W. L.; Tirado-Rives, J. The OPLS [Optimized Potentials for Liquid Simulations] Potential Functions for Proteins, Energy Minimizations for Crystals of Cyclic Peptides and Crambin. J. Am. Chem. Soc. 1988, 110 (6), 1657–1666.

(39) Van Zundert, G. C. P.; Rodrigues, J. P. G. L. M.; Trellet, M.; Schmitz, C.; Kastritis, P. L.; Karaca, E.; Melquiond, A. S. J.; Van Dijk, M.; De Vries, S. J.; Bonvin, A. M. J. J. The HADDOCK2.2 Web Server: User-Friendly Integrative Modeling of Biomolecular Complexes. J. Mol. Biol. 2016, 428 (4), 720–725.

(40) Jorgensen, W. L.; Tirado-Rives, J. The OPLS Potential Functions for Proteins. Energy Minimizations for Crystals of Cyclic Peptides and Crambin. J. Am. Chem. Soc. 1988, 110 (6), 1657–66.

(41) Rodrigues, J. P. G. L. M.; Trellet, M.; Schmitz, C.; Kastritis, P.; Karaca, E.; Melquiond, A. S. J.; Bonvin, A. M. J. J. Clustering Biomolecular Complexes by Residue Contacts Similarity. Proteins Struct. Funct. Bioinforma. 2012, 80 (7), 1810–7.

(42) Lensink, M. F.; Wodak, S. J. Docking and Scoring Protein Interactions: CAPRI 2009. Proteins Struct. Funct. Bioinforma. 2010, 78 (15), 3073–3084.

(43) Snijder, J.; Burnley, R. J.; Wiegard, A.; Melquiond, A. S. J.; Bonvin, A. M. J. J.; Axmann, I. M.; Heck, A. J. R. Insight into Cyanobacterial Circadian Timing from Structural Details of the KaiB-KaiC Interaction. Proc. Natl. Acad. Sci. 2014, pp 13008–13013.

(44) Tseng, R.; Goularte, N. F.; Chavan, A.; Luu, J.; Cohen, S. E.; Chang, Y. G.; Heisler, J.; Li, S.; Michael, A. K.; Tripathi, S.; et al. Structural Basis of the Day-Night Transition in a Bacterial Circadian Clock. Science (80-.). 2017, 355 (6330), 1174–1180.

(45) Villarreal, S. A.; Pattanayek, R.; Williams, D. R.; Mori, T.; Qin, X.; Johnson, C. H.; Egli, M.; Stewart, P. L. CryoEM and Molecular Dynamics of the Circadian KaiB-KaiC Complex Indicates That KaiB Monomers Interact with KaiC and Block ATP Binding Clefts. J. Mol. Biol. 2013, 425 (18), 3311–24.

(46) Lee, B.; Richards, F. M. The Interpretation of Protein Structures: Estimation of Static Accessibility. J. Mol. Biol. 1971, 55 (3), 379–400.

(47) Lensink, M. F.; Méndez, R.; Wodak, S. J. Docking and Scoring Protein Complexes: {CAPRI} 3rd {Edition}. Proteins Struct. Funct. Bioinforma. 2007, 69 (4), 704–18.

(48) Hayashi, F.; Suzuki, H.; Iwase, R.; Uzumaki, T.; Miyake, A.; Shen, J. R.; Imada, K.; Furukawa, Y.; Yonekura, K.; Namba, K.; et al. ATP-Induced Hexameric Ring Structure of the Cyanobacterial Circadian Clock Protein KaiC. Genes to Cells 2003, 8 (3), 287–296.

(49) Hayashi, F.; Iwase, R.; Uzumaki, T.; Ishiura, M. Hexamerization by the N-Terminal Domain and Intersubunit Phosphorylation by the C-Terminal Domain of Cyanobacterial Circadian Clock Protein KaiC. Biochem. Biophys. Res. Commun. 2006, 348 (3), 864–872.

(50) Snijder, J.; Schuller, J. M.; Wiegard, A.; Lössl, P.; Schmelling, N.; Axmann, I. M.; Plitzko, J. M.; Förster, F.; Heck, A. J. R. Structures of the Cyanobacterial Circadian Oscillator Frozen in a Fully Assembled State. Science (80-.). 2017, 355 (6330), 1181–1184.

(51) Pettersen, E. F.; Goddard, T. D.; Huang, C. C.; Couch, G. S.; Greenblatt, D. M.; Meng, E. C.; Ferrin, T. E. UCSF Chimera - A Visualization System for Exploratory Research and Analysis. J. Comput. Chem. 2004, 25 (13), 1605–12.

(52) Morin, A.; Eisenbraun, B.; Key, J.; Sanschagrin, P. C.; Timony, M. A.; Ottaviano, M.; Sliz, P. Collaboration Gets the Most out of Software. Elife 2013, 76, 821–828.

