## Supplementary Information for "Less is more: Coarse-grained integrative modeling of large biomolecular assemblies with HADDOCK"

### TABLE OF CONTENTS

**SI-1.** Implementation of the MARTINI force field for proteins in HADDOCK ............... 3

**Table SI-1.** Backbone particle type.

**Table SI-2.** Backbone-backbone relations.

**Table SI-3.** Side chains amino acid dependent and corresponding parameters.

**Table SI-4.** Polar and charged amino acid and corresponding parameters.

**SI-2.** Detailed overview of the selected cases from the Protein Docking Benchmark ......... 7

**Table SI-5.** Selected complexes for the CG protein-protein benchmark.

**SI-3.** Coarse-grained integrative modelling of KaiC-KaiB with HADDOCK .................. 8

**Table SI-6.** List of residues used as "active" to drive the docking.

**Table SI-7.** Paired i-RMSD after cross-superimposition of the two 6 fold rings in KaiC.

**Table SI-8.** Cluster based statistics for the CI and CII docking runs based on the fraction of common contacts.

**Table SI-9.** Structural similarity assessment of the top 4 models of coarse-grained HADDOCK with respect to the cryo-EM model.

**SI-4.** Reduction of the energy landscape ........................................................ 10

**Table SI-10.** Number of acceptable or higher quality models generated at the rigid-body (it0) stage of coarse-grained and standard all-atom HADDOCK (*ab-initio* mode)

**Table SI-11.** Number of acceptable or higher quality models generated at the rigid-body (it0) stage of coarse-grained and standard all-atom HADDOCK (*information-driven* mode)

#### SI-1. IMPLEMENTATION OF THE MARTINI FORCE FIELD FOR PROTEINS IN HADDOCK

Converted force field parameters from the original work of *de Jong. et al* into CNS compatible format.

**Table SI-1.** Backbone Particle Types.

| Amino Acid | Coil | Helix | Extended | Turn |
|---|---|---|---|---|
| **ALA** | F4 | HPa/HP0/HP5/Hda | B0 | T0 |
| **All others** | F5 | H0/H5/Hd/Ha | Bda | Tda |

**Table SI-2.** Backbone-Backbone Relations. D: Bond Length (Å). ϕ: Bond Angle (°).Ψ: Bond Dihedral (°). K: Force Constant (kcal.mol$^{-1}$).

| Backbone SS | $D_{BB}$ | $K_{BB}$ | $\phi_{BBB}$ | $K_{BBB}$ | $\Psi_{BBBB}$ | $K_{BBB}$ |
|---|---|---|---|---|---|---|
| **Coil** | 3.5 | 12.5 | 127 | 5.971 | - | - |
| **Helix** | 3.1 | 12.5 | 96* | 167.184 | -120 | 95.524 |
| **Extended** | 3.5 | 12.5 | 134 | 5.971 | 0 | 0.57 |
| **Turn** | 3.5 | 12.5 | 100 | 5.971 | - | - |

* $\phi_{BBB}$ = 98 ° and $K_{BBB}$ = 23.883 Kcal.mol$^{-1}$ for PRO in Helix conformation.

**Table SI-3.** Side-chains amino acid dependent and corresponding parameters. BB denotes Backbone-Backbone while BS Backbone-Side-chain.

| Amino Acid | SC Bead Name | $D_{BS}$,$D_{SS}$,$D_{SS}$,$D_{SS}$ | $K_{BS}$,$K_{SS}$,$K_{SS}$, $K_{SS}$ | $\phi_{BSS}$,$\phi_{BSS}$,$\phi_{SSS}$,$\phi_{SSS}$ | $K_{BSS}$,$K_{BSS}$,$K_{SSS}$,$K_{SSS}$ | $\Psi_{BSSS}$, $\Psi_{SSSS}$ | $K_{BSSS}$, $K_{SSSS}$ |
|---|---|---|---|---|---|---|---|
| **TRP** | WC4-SNd-SC5-SC5 | 3.00,2.70,2.70,2.70 | 50,500,500,500 | 210,90,50,50 | 11.942, 5.971,11.942,11.942 | 0, 0, 0 | 11.942,11.942,47.767 |
| **TYR** | YC4-SC4-SP1 | 3.20,2.70, - | 50,500,500, - | 150,150, - | 11.942,11.942, - | 0, - | 11.942, - |
| **PHE** | FC5-SC5-SC5 | 3.10,2.70, - | 75,500,500, - | 150,150, - | 11.942,11.942, - | 0, - | 11.942, - |
| **HIS** | HC4-SP1-SP1 | 3.20,2.70, - | 75,500,500, - | 150,150, - | 11.942,11.942, - | 0, - | 11.942, - |
| **CYS** | C5 | 3.10, - | 75, - | - | - | - | - |
| **ILE** | AC1 | 3.10, - | 12.5, - | - | - | - | - |
| **LEU** | LC1 | 3.30, - | 75, - | - | - | - | - |
| **MET** | MC5 | 4.00, - | 25, - | - | - | - | - |
| **PRO** | PC3 | 3.00, - | 75, - | - | - | - | - |
| **VAL** | AC2 | 2.65, - | 12.5, - | - | - | - | - |
| **ALA** | - | - | - | - | - | - | - |
| **GLY** | - | - | - | - | - | - | - |

D: Bond Length (Å). ϕ: Bond Angle (°).Ψ: Bond Dihedrals and Improper (°).

**Table SI-4.** Polar and charged amino acid and corresponding parameters beads including "fake-beads" for a better description of electrostatics. BB denotes Backbone-Backbone while BS Backbone-Side-chain. Q: Charge (electron charge units). D: Bond Length (Å). ϕ: Bond Angle (°).

| Amino Acid | SC Bead Name | Q (SCd) | $D_{BS}$,$D_{SS}$,$D_{SS}$ | $K_{BS}$,$K_{SS}$,$K_{SS}$ | $\phi_{BSS}$ | $K_{BSS}$ |
|---|---|---|---|---|---|---|
| **ARG** | RN0-AQd-SCd | +1.00 | 3.30,3.4,1.10 | 50,50,500 | 180 | 5.971 |
| **LYS** | KC3-KQd-SCd | +1.00 | 3.30,2.80,1.10 | 50,50,500 | 180 | 5.971 |
| **ASN** | Nda-SCd-SCd | ±0.46 | 3.20,1.10,2.80 | 50,500,500 | - | - |
| **GLN** | QNda-SCd-SCd | ±0.46 | 4.00,1.10,2.80 | 50,500,500 | - | - |
| **SER** | SN0-SCd-SCd | ±0.40 | 2.50,1.10,2.80 | 75,500,500 | - | - |
| **THR** | TNda-SCd-SCd | ±0.31 | 2.50,1.10,2.80 | 75,500,500 | - | - |
| **GLU** | Qa-SCd | -1.00 | 4.00,1.10 | 50,500 | - | - |
| **ASP** | DQa-SCd | -1.00 | 3.20,1.10 | 75,500 | - | - |

#### SI-2. DETAILED OVERVIEW OF THE SELECTED CASES FROM THE PROTEIN DOCKING BENCHMARK 5, N = 27

**Table SI-5**. Selected complexes (27) for the CG protein-protein benchmark. Classified in the Protein-Protein Benchmark version 5.0 as: enzyme-inhibitor, enzyme-substrate, enzyme complex or others.

| Complex | # Atoms | # Residues | MW | Resolution | Receptor | Resolution | Ligand | Resolution |
|---|---|---|---|---|---|---|---|---|
| **1AZS** Adenylyl cyclase – GTPgammaS | 5740 | 718 | 77020 | 2,3 | **1AB8** | 2,2 | **1AZT** | 2,3 |
| **1DE4** HFE – Transferrin R | 13107 | 1641 | 172501 | 2,8 | **1A6Z** | 2,6 | **1CX8** | 3,2 |
| **1EXB** T1 β – $K^+$ channel | 13256 | 1668 | 177850 | 2,1 | **1QRQ** | 2,8 | **1QDV** | 1,6 |
| **1GP2** Protein G trimer | 5781 | 737 | 77145 | 2,3 | **1GIA** | 2 | **1TBG** | 2,5 |
| **1GXD** proMMP2 – TIMP2 | 6466 | 816 | 85521 | 3,1 | **1CK7** | 2,8 | **1BR9** | 2,1 |
| **1H1V** Actin – Gelsolin | 5414 | 695 | 71583 | 2,99 | **1IJJ** | 2,85 | **1D0N** | 2,5 |
| **1HE8** Ras – PI3 γ kinase | 7396 | 915 | 98167 | 3 | **821P** | 1,5 | **1E8Z** | 2,4 |
| **1IB1** 14-3-3 protein – N-acetylase | 5046 | 632 | 70414 | 2,7 | **1QJB** | 2 | **1KUY** | 2,4 |
| **1KXP** Actin – Vitamin D | 6167 | 787 | 82409 | 2,1 | **1IJJ** | 2,85 | **1KW2** | 2,15 |
| **1N2C** Nitrogenase | 20058 | 2548 | 265698 | 3 | **3MIN** | 2,03 | **2NIP** | 2,2 |
| **1RLB** Transthyretin – Retinol | 5171 | 660 | 68709 | 3,1 | **2PAB** | 1,8 | **1HBP** | 1,9 |
| **1T6B** Anthrax – Anthrax receptor | 6695 | 846 | 88168 | 2,5 | **1ACC** | 2,1 | **1SHU** | 1,5 |
| **1WDW** Tryptophan synthase | 7848 | 1011 | 103279 | 3 | **1V8Z** | 2,21 | **1GEQ** | 2 |
| **1Y64** Actin – BNI1 | 6119 | 767 | 81284 | 3 | **2FXU** | 1,35 | **1UX5** | 2,5 |
| **2AJF** ACE2 – SARS | 6273 | 771 | 82831 | 2,9 | **1R42** | 2,2 | **2GHV** | 2,2 |
| **2FJU** Phospolipase β2 – Rac | 6993 | 873 | 92947 | 2,2 | **2ZKM** | 1,62 | **1MH1** | 1,38 |
| **2GAF** VP55 – VP39 | 5929 | 723 | 79736 | 2,4 | **3OWG** | 2,86 | **1VPT** | 1,8 |
| **2OOR** NAD(P) α – NAD(P) β | 6755 | 915 | 90178 | 2,32 | **1L7E** | 1,9 | **1E3T** | NMR |
| **3AAA** Actin – Myotrophin | 5033 | 634 | 66478 | 2,2 | **3AA7** | 1,9 | **1MYO** | NMR |
| **3BIW** Neuroglin | 5545 | 710 | 72968 | 3,5 | **3BIX** | 2,61 | **2R1D** | 2,6 |
| **3L89** Ad21 – CD46 | 5252 | 677 | 69446 | 3,5 | **3L88** | 2,5 | **1CKL** | 3,1 |
| **3LVK** IscS – tusA | 6641 | 850 | 88578 | 2,442 | **3LVM** | 2,33 | **1DCJ** | NMR |
| **3R9A** Alanine-glyoxilate AT – PEX5P | 8199 | 1063 | 108889 | 2,35 | **1H0C** | 2,5 | **2C0M** | 2,5 |
| **4GAM** Methane monooxygenase | 18499 | 2262 | 243436 | 2,902 | **1XVB** | 1,8 | **1CKV** | NMR |
| **4H03** IA – Actin α | 6152 | 770 | 82388 | 1,75 | **1GIQ** | 1,8 | **1IJJ** | 2,85 |
| **4JCV** RecR – RecO | 7529 | 993 | 99383 | 3,34 | **1VDD** | 2,5 | **1W3S** | 2,4 |
| **4LW4** CsdA – CsdE | 7095 | 935 | 93969 | 2,01 | **4LW2** | 1,8 | **1NI7** | NMR |

#### SI-3. COARSE-GRAINED INTEGRATIVE MODELLING of KaiC-KaiB WITH HADDOCK

**Table SI-6.** Detailed list of residues, identified by mutagenesis experiments in combination with hydrogen-deuterium exchange and mass spectroscopy (HDX-MS), used as "active" in HADDOCK-CG to drive the simulations (CI/CII).

| Protein | Domain | Residues |
|---|---|---|
| KaiC | C I | Gly101, Leu103, Ile105, Leu106, Asp107, Ala108, Pro110, Asp111, Pro112, Glu113, Gly114, Gln115, Glu116, Val117, Val118, Gly119, Asp122, Leu123, Ser124, Ala125, Leu126, Ile130, Ala133, Ile134 |
| | C II | Met449, Ser450, Arg451, Ala452, Ile453, Asn454, Val455, Phe456, Lys457, Met458, Arg459, Gly460, His463, Asp464, Lys465, Ala466, Ile467, Arg468, Glu469, Phe470 |
| KaiB | | Thr7, Asn17, Thr18, Pro19, Glu33, Glu35, Gly38, Lys43, Leu48, Lys49, Pro51, Gln52, Glu55, Glu56, Lys58, Leu60, Pro70, Pro71, Pro72, Val73, Arg74, Ile77, Ser81, Asn82, Glu84, Lys85, Ile88 |

**Table SI-7**. Paired i-RMSD values (Å) calculated after cross-superimposition of the two 6 fold rings in KaiC.

| KaiC | A - 1 | A - 2 | A - 3 | A - 4 | A - 5 | A - 6 |
|---|---|---|---|---|---|---|
| A - 1 | - | 0.96 | 1.29 | 1.9 | 0.8 | 0.89 |
| A - 2 | 0.96 | - | 1.23 | 1.89 | 0.91 | 1.01 |
| A - 3 | 1.29 | 1.23 | - | 1.28 | 0.89 | 1.25 |
| A - 4 | 1.9 | 1.89 | 1.28 | - | 1.4 | 1.39 |
| A - 5 | 0.8 | 0.91 | 0.89 | 1.4 | - | 0.92 |
| A - 6 | 0.89 | 1.01 | 1.25 | 1.39 | 0.92 | - |

**Table SI-8.** Cluster based statistics for the CI and CII docking runs based on the fraction of common contacts (0.5 cutoff). HADDOCK score single terms averaged over the top 4 members of each cluster are reported and clusters are ordered according to the averaged HADDOCK score (a.u.). $E_{vdw}$: Lennard-Jones potential. $E_{elec}$: Coulomb potential. $E_{AIR}$: Ambiguous interaction restraints energy. $E_{desolv}$: Empirical desolvation score. BSA: Buried surface area. $E_{symmetry}$: Symmetry restraints energy.

| Cluster | Population | $E_{vdw}$ | $E_{elec}$ | $E_{AIR}$ | $E_{desolv}$ | BSA | $E_{symmetry}$ | HADDOCK score | I-RMSD [Å] |
|---|---|---|---|---|---|---|---|---|---|
| **CI domain** | | | | | | | | | |
| **1** | 15 | -366.3 ± 28.4 | -809.3 ± 153 | 2899.2 ± 126.6 | 12.3 ± 12.3 | 11598.3 ± 799.5 | 91.5 ± 36.8 | **-216.7 ± 13.2** | 5.9 ± 1.3 |
| **3** | 9 | -343.6 ± 18.1 | -917.3 ± 178.2 | 2946 ± 103.3 | 29 ± 19.1 | 10805.3 ± 611 | 114.2 ± 53.7 | **-191.9 ± 30.5** | 21.1 ± 6.6 |
| **9** | 4 | -327.8 ± 18.7 | -1038.3 ± 145 | 3076.3 ± 350.6 | 27.6 ± 32.8 | 10905.4 ± 391.4 | 88.2 ± 19.6 | **-191.3 ± 43** | 18.3 ± 6.5 |
| **2** | 10 | -329.3 ± 16.8 | -1033.3 ± 122.7 | 3303.7 ± 231 | 36.3 ± 20 | 10669.7 ± 890.1 | 134.9 ± 63.5 | **-160.3 ± 16.9** | 19 ± 5.3 |
| **6** | 4 | -304.9 ± 39.4 | -897.2 ± 116.6 | 3154.9 ± 250.8 | 6.9 ± 18.2 | 10432.8 ± 996.9 | 78.2 ± 5.5 | **-154.1 ± 52.6** | 16.8 ± 2.9 |
| **CII domain** | | | | | | | | | |
| **2** | 4 | 2206.6 ± 214.5 | -265 ± 27.4 | 21375.2 ± 5169.3 | -28.3 ± 77.8 | 14969.5 ± 5869.1 | 156.3 ± 45.48 | **+44.5 ± 19** | - |

**Table SI-9.** Structural similarity assessment of the top 4 models of coarse-grained HADDOCK (from the best cluster) with respect to the cryo-EM (backbone only) model (PDB ID: 5N8Y). B/C/D/E/F/G correspond to the 6 KaiB monomers, respectively, docked onto KaiC. i-RMSD, l-RMSD and FNAT are calculated according to CAPRI criteria.

| KaiB Subunits | i-RMSD [Å] | l-RMSD [Å] | FNAT |
|---|---|---|---|
| **Overall** | **10.1 ± 2.8** | **5.9 ± 1.3** | **0.09 ± 0.05** |
| B | 8.4 ± 2.2 | 3.3 ± 0.4 | 0.07 ± 0.07 |
| C | 8.3 ± 2.4 | 3.4 ± 0.4 | 0.05 ± 0.06 |
| D | 8.8 ± 2.1 | 3.5 ± 0.4 | 0.05 ± 0.07 |
| E | 8.3 ± 2.2 | 3.4 ± 0.4 | 0.11 ± 0.08 |
| F | 8.4 ± 2.0 | 3.3 ± 0.4 | 0.05 ± 0.05 |
| G | 6.7 ± 2.8 | 3.3 ± 0.4 | 0.12 ± 0.06 |

#### SI-4. REDUCTION OF THE ENERGY LANDSCAPE COMPLEXITY

**Table SI-10**. Number of acceptable or higher quality models, for each of the protein docking benchmark complexes, generated at the rigid-body (it0) stage of coarse-grained and standard all-atom HADDOCK docking runs in the absence of information to drive the docking (*ab-initio* mode). 10000 models were generated in the case of *ab-initio* docking.

| Complex | Protocol | Top 200 | Top 400 | Total |
|---|---|---|---|---|
| 1AZS | Coarse-grained | 0 | 0 | 4 |
| | All-atom | 0 | 0 | 1 |
| 1DE4 | Coarse-grained | 0 | 0 | 4 |
| | All-atom | 0 | 0 | 0 |
| 1EXB | Coarse-grained | 0 | 0 | 0 |
| | All-atom | 0 | 0 | 0 |
| 1GP2 | Coarse-grained | 0 | 1 | 4 |
| | All-atom | 2 | 3 | 3 |
| 1GXD | Coarse-grained | 1 | 1 | 2 |
| | All-atom | 1 | 1 | 1 |
| 1H1V | Coarse-grained | 0 | 0 | 0 |
| | All-atom | 0 | 0 | 0 |
| 1HE8 | Coarse-grained | 4 | 4 | 4 |
| | All-atom | 0 | 0 | 6 |
| 1IB1 | Coarse-grained | 0 | 0 | 0 |
| | All-atom | 0 | 0 | 0 |
| 1KXP | Coarse-grained | 2 | 2 | 2 |
| | All-atom | 0 | 0 | 2 |
| 1N2C | Coarse-grained | 0 | 0 | 0 |
| | All-atom | 0 | 0 | 0 |
| 1RLB | Coarse-grained | 0 | 0 | 3 |
| | All-atom | 0 | 0 | 0 |
| 1T6B | Coarse-grained | 2 | 2 | 17 |
| | All-atom | 0 | 0 | 19 |
| 1WDW | Coarse-grained | 0 | 0 | 4 |
| | All-atom | 3 | 3 | 3 |
| 1Y64 | Coarse-grained | 0 | 0 | 0 |
| | All-atom | 0 | 0 | 0 |
| 2AJF | Coarse-grained | 0 | 0 | 1 |
| | All-atom | 0 | 0 | 0 |
| 2FJU | Coarse-grained | 0 | 0 | 4 |
| | All-atom | 0 | 0 | 1 |
| 2GAF | Coarse-grained | 0 | 0 | 0 |
| | All-atom | 0 | 0 | 2 |
| 2OOR | Coarse-grained | 0 | 0 | 0 |
| | All-atom | 0 | 0 | 0 |
| 3AAA | Coarse-grained | 0 | 0 | 1 |
| | All-atom | 0 | 0 | 0 |
| 3BIW | Coarse-grained | 0 | 0 | 5 |
| | All-atom | 0 | 0 | 5 |
| 3L89 | Coarse-grained | 1 | 1 | 1 |
| | All-atom | 0 | 0 | 2 |
| 3LVK | Coarse-grained | 2 | 2 | 5 |
| | All-atom | 1 | 1 | 2 |
| 3R9A | Coarse-grained | 0 | 0 | 0 |
| | All-atom | 0 | 0 | 0 |
| 4GAM | Coarse-grained | 0 | 0 | 0 |
| | All-atom | 0 | 0 | 0 |
| 4H03 | Coarse-grained | 2 | 2 | 7 |
| | All-atom | 3 | 4 | 4 |
| 4JCV | Coarse-grained | 1 | 1 | 2 |
| | All-atom | 0 | 0 | 1 |
| 4LW4 | Coarse-grained | 0 | 0 | 4 |
| | All-atom | 1 | 1 | 1 |
| TOTAL | **Coarse-grained** | **15** | **16** | **74** |
| | **All-atom** | **11** | **13** | **53** |

**Table SI-11**. Number of acceptable or higher quality models, for each of the protein docking benchmark complexes, generated at the rigid-body (it0) stage of the coarse-grained and standard all-atom HADDOCK docking runs using true interface information to drive the docking. 1000 models were generated in the case of *information-driven* docking.

| Complex | Protocol | Top 200 | Top 400 | Total |
|---|---|---|---|---|
| 1AZS | Coarse-grained | 170 | 270 | 332 |
| | All-atom | 65 | 117 | 213 |
| 1DE4 | Coarse-grained | 2 | 6 | 120 |
| | All-atom | 137 | 264 | 353 |
| 1EXB | Coarse-grained | 36 | 73 | 143 |
| | All-atom | 33 | 69 | 185 |
| 1GP2 | Coarse-grained | 111 | 148 | 155 |
| | All-atom | 152 | 206 | 215 |
| 1GXD | Coarse-grained | 102 | 148 | 184 |
| | All-atom | 66 | 118 | 146 |
| 1H1V | Coarse-grained | 0 | 0 | 0 |
| | All-atom | 8 | 26 | 48 |
| 1HE8 | Coarse-grained | 157 | 285 | 491 |
| | All-atom | 53 | 63 | 212 |
| 1IB1 | Coarse-grained | 0 | 0 | 0 |
| | All-atom | 1 | 1 | 9 |
| 1KXP | Coarse-grained | 199 | 395 | 563 |
| | All-atom | 188 | 337 | 511 |
| 1N2C | Coarse-grained | 0 | 1 | 4 |
| | All-atom | 34 | 70 | 183 |
| 1RLB | Coarse-grained | 160 | 320 | 618 |
| | All-atom | 190 | 357 | 734 |
| 1T6B | Coarse-grained | 184 | 367 | 803 |
| | All-atom | 143 | 324 | 667 |
| 1WDW | Coarse-grained | 200 | 397 | 680 |
| | All-atom | 200 | 399 | 620 |
| 1Y64 | Coarse-grained | 0 | 0 | 0 |
| | All-atom | 0 | 0 | 0 |
| 2AJF | Coarse-grained | 66 | 156 | 412 |
| | All-atom | 111 | 257 | 422 |
| 2FJU | Coarse-grained | 126 | 253 | 694 |
| | All-atom | 165 | 347 | 703 |
| 2GAF | Coarse-grained | 131 | 250 | 573 |
| | All-atom | 157 | 266 | 570 |
| 2OOR | Coarse-grained | 17 | 18 | 25 |
| | All-atom | 38 | 43 | 46 |
| 3AAA | Coarse-grained | 52 | 132 | 289 |
| | All-atom | 128 | 230 | 369 |
| 3BIW | Coarse-grained | 107 | 245 | 627 |
| | All-atom | 3 | 21 | 218 |
| 3L89 | Coarse-grained | 109 | 230 | 488 |
| | All-atom | 132 | 253 | 430 |
| 3LVK | Coarse-grained | 199 | 386 | 708 |
| | All-atom | 188 | 301 | 411 |
| 3R9A | Coarse-grained | 152 | 327 | 790 |
| | All-atom | 113 | 260 | 725 |
| 4GAM | Coarse-grained | 0 | 0 | 0 |
| | All-atom | 0 | 0 | 0 |
| 4H03 | Coarse-grained | 186 | 376 | 606 |
| | All-atom | 90 | 215 | 454 |
| 4JCV | Coarse-grained | 173 | 250 | 289 |
| | All-atom | 198 | 252 | 266 |
| 4LW4 | Coarse-grained | 27 | 33 | 95 |
| | All-atom | 109 | 144 | 186 |
| TOTAL | **Coarse-grained** | **2666** | **5066** | **9689** |
| | **All-atom** | **2702** | **4940** | **8896** |
